# First-in-patient dose prediction for adeno-associated virus-mediated hemophilia gene therapy using allometric scaling

**DOI:** 10.1101/2022.07.05.498837

**Authors:** Peng Zou

## Abstract

Rational first-in-patient (FIP) dose selection is critical for successful clinical development of gene therapy. In this study, the author compared the performance of two allometric scaling approaches and body weight-based dose conversion approach for FIP dose prediction for adeno-associated virus (AAV)-mediated hemophilia gene therapy. The performance of the three approaches was examined using preclinical and clinical efficacy data of nine AAV vectors. In general, body weight-based direct conversion of effective doses in monkeys or dogs was more likely to underestimate FIP dose but worked better for one bioengineered vector with a high transduction efficiency specifically in humans. In contrast, allometric scaling between gene efficiency factor (logGEF) and body weight (logW) was likely to overestimate FIP dose but worked better than the other two approaches when vector capsid-specific T-cell responses were detected in patients. The third approach, allometric scaling between logGEF and W^-0.25^ was appropriate for FIP dose prediction in the absence of T-cell responses to AAV vectors or a dramatic difference in vector transduction efficiency between animals and humans. A decision tree was tentatively proposed to facilitate approach selection for FIP dose prediction in AAV-mediated hemophilia gene therapy.

## Introduction

Rational first-in-patient (FIP) dose selection is critical for successful clinical development of gene therapy, especially for rare disease programs. The target population of many gene therapy products is a small population with rare genetic diseases. The safety risks of most gene therapy products include the possibility of extended or permanent effects ^1^. Therefore, the benefit-risk profile is not acceptable for healthy volunteers in many gene therapy clinical trials ^1^. The first-in-human study of many gene therapy products is a phase 1/2 dose-finding study conducted in a dozen or so patients with a rare disease. Each patient enrolled in the dose-finding study often receives a single dose of adeno-associated virus (AAV) vector only because the immune responses caused by repeated dosing may render subsequent administrations ineffective. Especially for patients with serious or life-threatening genetic diseases, it is not feasible or ethical to use a low and ineffective starting dose, which may exclude that patient from any future benefit of this treatment. The FIP dose is anticipated to provide therapeutic benefit to patients. On the other hand, a high-dose AAV gene therapy may cause severe adverse events even deaths. Two X-linked myotubular myopathy patients with pre-existing liver disease died from liver dysfunction after they received a high dose (3 × 10^14^ vg/kg) of adeno-associated virus-8 (AAV8) vector-based investigational gene therapy (AT132) ^2^. Therefore, an accurate prediction of FIP dose of gene therapy products is critical. Compared to small molecules and therapeutic proteins, our knowledge of FIP dose prediction for gene therapy is still limited. Traditional quantitative methods such as PK-PD modeling used to predict human dose of small molecules and therapeutic proteins are difficult to apply to gene therapy, model structures and parameter paradigms may not be directly applicable to these complex gene therapy products ^3^.

Due to its relatively low genotoxicity profile in human, high titer, mild immune response, and ability to infect a broad range of cells, recombinant AAV is a preferred vector for delivering genes to target cells ^4^. Both the efficacy and safety of AAV-based gene therapy are largely dose-dependent ^5^. Rational prediction of FIP dose is critical for a dose-finding study of AAV-mediate gene therapy. Currently, a direct vg/kg conversion of no-observed-adverse-effect level (NOAEL) dose or pharmacologically active dose (PAD) from animal species is commonly used to predict FIP dose of systemically administered AAV-mediated gene therapy and an empirical scaling factor is usually applied to further reduce the starting dose^6^. However, the direct vg/kg conversion approach is likely to provide an ineffective FIP dose in AAV-mediated hemophilia B therapy ^7^.

Recent investigations in allometric scaling-based dose prediction have shed light on FIP dose selection for AAV-mediated gene therapy ^7, 8^. It was reported that the metabolic rate of cells (B_c_) in vivo in mammalian species decreased with increasing body weight (B_c_ = B_0_ * W^-0.25^, where B_0_ is the cellular metabolic rate expressed as 3 × 10^-11^ Watts per cell for organism with mass of 1.0 gram, and W is the mass of a mammalian species) ^9^. Since intracellular synthesis of the protein of interest following gene delivery is dependent on metabolic rate of cells, Tang et. al. introduced a concept of gene efficiency factor (GEF) to describe the efficiency of the gene transfer system and demonstrated a linear relationship between logGEF and logW using preclinical and clinical data of three AAV vectors for hemophilia B therapy, where W was the body weight of mammalian species^7^. Tang et. al. concluded that body weight-based allometric scaling of GEF could be used to predict FIP dose of AAV-mediated gene therapy. However, according to the relationship between intracellular protein synthesis and W^-0.25^, this author believes it is more appropriate to use W^-0.25^ instead of logW as the variant for allometric scaling of logGEF.

Hemophilia, an inherited bleeding disorder, was one of the earliest diseases considered for gene therapy. The most common types of hemophilia are hemophilia A and hemophilia B, caused by mutations in *F8* or *F9*, coding for factor VIII (FVIII) and factor IX (FIX) proteins, respectively ^10^. FIP dose selection for gene therapy should be based on the targeted plasma levels FVIII and FIX. Natural history data from hemophilia A patients indicate that endogenous plasma factor VIII activity of 12% (12 IU/dL) or more of the normal value in healthy subjects eliminates spontaneous hemarthrosis ^11^. Patients with mild hemophilia B, who have plasma FIX level of 5% - 40% of normal value, may appear phenotypically normal and never show signs of uncontrolled bleeds unless undergoing severe trauma or surgery. Therefore, it appears reasonable to propose 12% and 5% of normal plasma levels as the targeted FVIII and FIX levels, respectively, for FIP dose selection. In this study, the performance of three approaches for FIP dose predictions, including allometric scaling with logW (AS-logW), allometric scaling with W^-0.25^ (AS-W^-0.25^) and direct vg/kg conversion from preclinical dose, was compared using preclinical and clinical data of nine AAV vectors.

## Methods

### Data source

The preclinical and clinical targeted protein expression data of vectors were collected from published literature. Only the vectors with clinical data and preclinical data from ≥ 2 species available were collected. When the body weight of individual subject or animal was not available in literature reports, the body weights of mouse, cynomolgus macaque, rhesus macaque, and human were assumed as 0.02, 2.5, 8.0, and 70 kg, respectively.

### GEF calculation and allometric scaling approaches

Individual animal and human GEF values were calculated using previously reported Equation 1 ^7^.

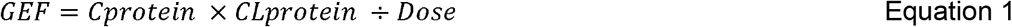

C_protein_ was the steady-state maximum plasma concentration (C_max_) of transgene product following gene therapy. CL_protein_ was the total clearance of FIX or FVIII, which was the average clearance values from multiple animal/human pharmacokinetic studies (Table S1). Dose was the total vector genomes (vg) received by individual animal or patient. The mean GEF of each species was used for allometric scaling. For the allometric scaling approaches, regression analyses were conducted between logGEF and logW or W^-0.25^ using the data of three species (two animal species and human) and two animal species only. The equation derived from two animal species was used to predict human GEF.

### Targeted plasma transgene product levels and FIP dose projection

The FIP dose is anticipated to be effective in eliminating uncontrolled bleeds. Thus, 12% of normal plasma FVIII activity level and 5% of normal plasma FIX level were proposed as the targeted protein levels for hemophilia A and B gene therapy, respectively. The normal level of FIX in the plasma of healthy subjects is 5000 ng/mL ^12^ and thus a targeted FIX level of 250 ng/mL was used to project FIP doses of FIX vectors except for SPK-9001. FIX-Padua is a hyperfunctional factor IX and the residue 338 in human FIX is changed from arginine to alanine. FIX-Pauda expressed by SPK-9001 has an 8- to 12-fold increased specific activity (a mean difference of 9.1-fold in hemophilia B mice) compared to endogenous FIX ^13^. Therefore, the targeted plasma FIX-Pauda level was selected as 250 ng/mL ÷ 9.1 = 27.5 ng/mL (equivalent to 5% of normal level). Equation 2 was used to predict FIP dose.

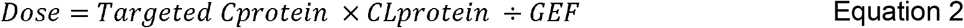

Where the targeted C_protein_ was 12 IU/dL for FVIII and 250 ng/mL for FIX (27.5 ng/mL for SPK-9001). GEF is the predicted human GEF derived from allometric scaling of two animal species.

### Direct vg/kg conversion approach

A linear regression was conducted between monkey or dog plasma FIX levels or FVIII activity levels at various doses and total vector doses (vg). Then, plasma FIX level of 250 ng/mL (27.5 ng/mL for SPK-9001) or plasma FVIII activity level of 12 IU/dL was incorporated into the regression equation to achieve monkey or dog total vector dose which was anticipated to produce targeted level of transgene product in monkeys or dogs. The total dose was divided by animal body weight and the weight normalized dose (vg/kg) is the FIP dose derived from the direct vg/kg conversion approach.

## Results

A total of five vectors for hemophilia A gene therapy and six vectors for hemophilia B gene therapy were identified (Supplementary materials, Tables S1 and S2). For vectors AMT-060 and AMT061 for hemophilia B gene therapy, only plasma FIX data from monkeys and humans were available (Table S2). The GEF values were calculated but allometric scaling could not be conducted. Thus, the data of five FVIII vectors and four FIX vectors were applied to allometric scaling analyses. In the dataset of rAAV2-hAAT-FIX vector, Tang et. al., excluded Subject F because the patient’s plasma FIX quickly decreased from peak level (150 ng/mL or 3% of normal level) following the infusion of rAAV2-hAAT-FIX vector due to T-cell responses to the vector ^7, 14, 15^. Both subjects (Subjects E and F) receiving 2×10^12^ vg/kg rAAV2-hAAT-FIX vector exhibited T-cell response against capsid ^14, 15^. To assess the impact of T-cell responses on the performance of allometric scaling approaches, both subjects were included in our analysis.

The preclinical and clinical doses, plasma FVIII activity or FIX levels, and calculated GEF for the 11 vectors were summarized in Tables S1 and S2. Using the mean clearance values of FIX and FVIII proteins (Table S3), individual subject’s or animal GEF value was derived from observed plasma protein level and vector dose. Consistent with previous studies ^7, 8^, a negative linear relationship between logGEF and logW of two preclinical species and humans was observed for the nine vectors (Figures 1 and 2). LogGEF decreases with increasing logW. Not surprisingly, a positive linear relationship between logGEF and W^-0.25^ was observed for the nine vectors (Figures 1 and 2). AS-W^-0.25^ generated a higher correlation coefficient (R^2^) than AS-logW for 6 among 9 vectors. The lowest R^2^ value of AS-W^-0.25^ and AS-logW based correlations was 0.88 (SB-525 vector) and 0.78 (SPK-8011 vector), respectively. The results of allometric scaling analyses suggest that gene expression efficiency is reversely corelated with body weight of mammalian species.

**Figure 1.**
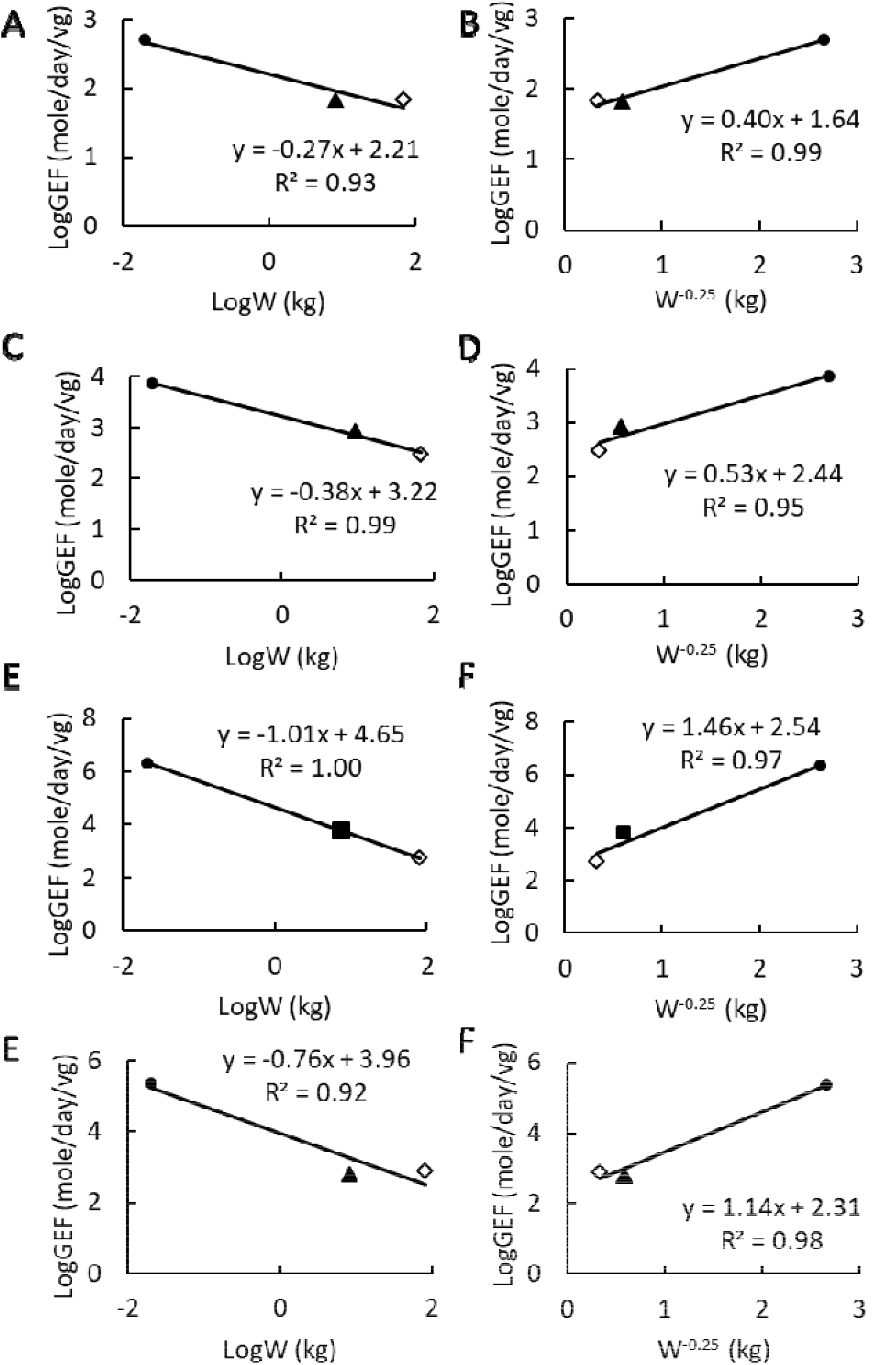
Allometric scaling of gene efficiency factor (GEF) for (A and B) rAAV2-CMV-FIX, (C and D) rAAV2-hAAT-FIX, (E and F) scAAV2/8-LP1-hFIXco, and (G and H) rAAV-Spark100-FIX-R338L. LogW was used as the scaling variable in A, C, E, and G and W^-0.25^ was used as the scaling variable in B, D, F, and H. W is body weight in kg and the unit of GEF is molecules/day/viral genome. Circle, triangle, square, and diamond symbols represent mouse, dog, macaque, and human, respectively.

**Figure 2.**
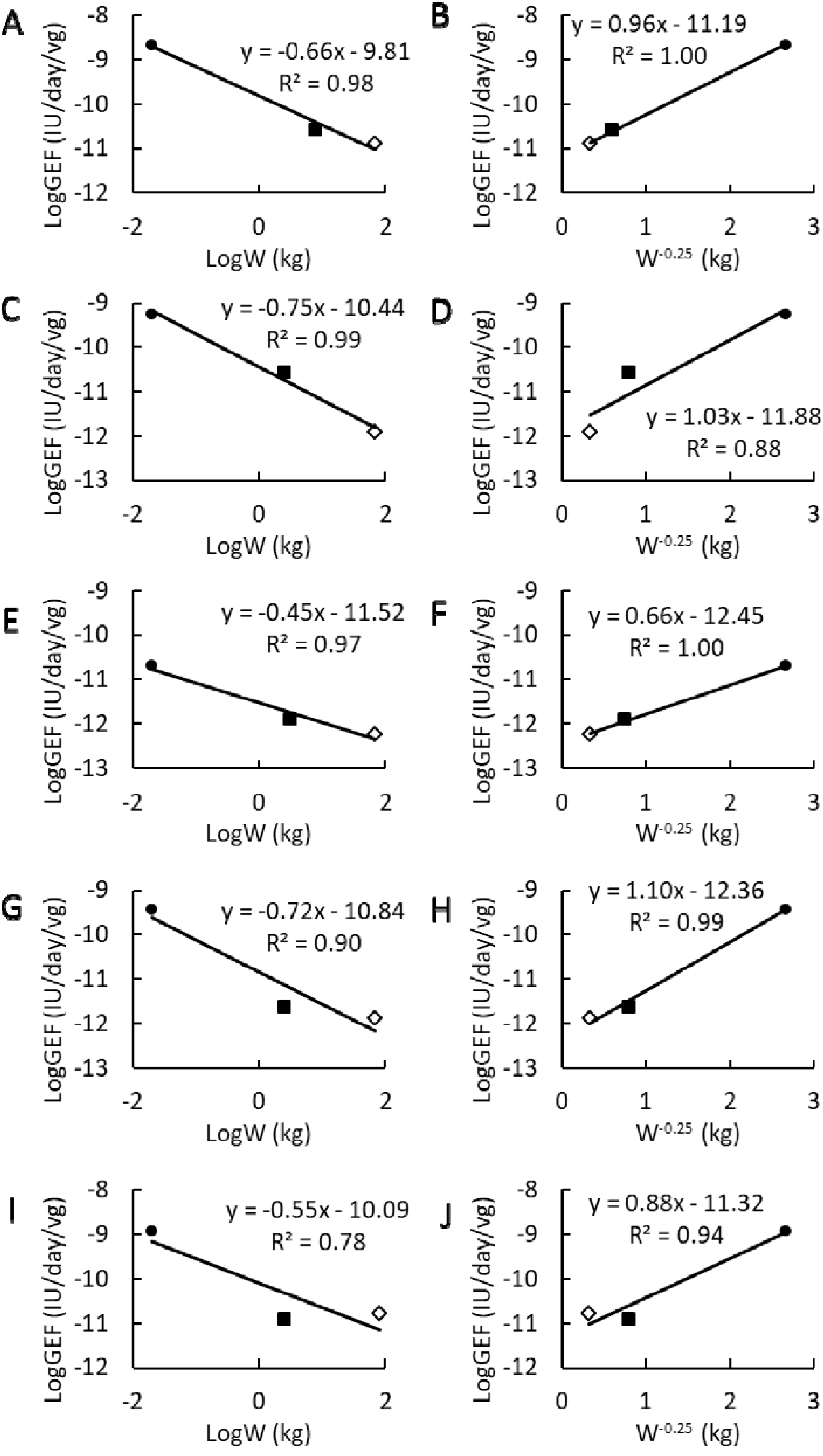
Allometric scaling of gene efficiency factor (GEF) for (A and B) GO-8, (C and D) SB-525, (E and F) BMN270, (G and H) DTX201, and (I and J) SPK-8011. LogW was used as the scaling variable in A, C, E, G, and I and W^-0.25^ was used as the scaling variable in B, D, F, H, and J. W is body weight in kg and the unit of GEF is IU/day/viral genome. Circle, square, and diamond symbols represent mouse, macaque, and human, respectively.

To predict human mean GEF, the preclinical data from two species were applied to logW- and W^-0.25^-dependent allometric scaling for the 9 vectors. The plots and derived linear equations were summarized in Figures S1 and S2 and Tables S4 and S5. The predicted Human GEF values derived from the two allometric scaling approaches were summarized in Table 1. For the 9 predicted human GEF values, 2 among 9 predictions by AS-logW were within 2-fold errors and 4 among 9 predictions by AS-W^-0.25^ were within 2-fold errors. AS-logW caused 39-fold underprediction of human GEF for SPK-8011 while AS-W^-0.25^ resulted in 11-fold overprediction of human GEF for SB-525.

**Table 1.** A comparison of human gene efficiency factor (GEF) predicted using two allometric scaling approaches versus observed human GEF

| Gene therapy | | Allometric scaling<br>using logW | | Allometric scaling<br>using $W^{-0.25}$ | | Observed<br>human<br>mean GEF |
| --- | --- | --- | --- | --- | --- | --- |
|  |  | Predicted<br>human<br>GEF | Error<br>(fold) | Predicted<br>human GEF | Error<br>(fold) |  |
| Factor IX<br>(molecules/day/vg) | rAAV2-CMV-FIX | 31.3 | ↓2.13 | 51.1 | ↓1.31 | 66.8 |
|  | rAAV2-hAAT-FIX | 414 | ↑1.40 | 661 | ↑2.23 | 296 |
|  | scAAV2/8-LP1-hFIXco | 557 | ↑1.07 | 2745 | ↑5.28 | 520 |
|  | rAAV-Spark100-FIX-R338L (SPK-9001) | 65 | ↓11.3 | 296 | ↓2.29 | 737 |
| Factor VIII<br>(IU/day/vg) | rAAV8-HLP-hFVIII-V3 (GO-8) | $5.11 \times 10^{-12}$ | ↓2.54 | $1.50 \times 10^{-11}$ | ↑1.16 | $1.30 \times 10^{-11}$ |
| | Giroctocogene fitelparvovec (SB-525) | $3.33 \times 10^{-12}$ | ↑2.79 | $1.31 \times 10^{-11}$ | ↑11.0 | $1.19 \times 10^{-12}$ |
| | Valoctocogene roxaparvovec (BMN270) | $2.14 \times 10^{-13}$ | ↓2.67 | $6.64 \times 10^{-13}$ | ↑1.16 | $5.72 \times 10^{-13}$ |
| | Peboctocogene camaparvovec (DTX201) | $7.11 \times 10^{-14}$ | ↓18.6 | $7.00 \times 10^{-13}$ | ↓1.89 | $1.33 \times 10^{-12}$ |
| | Dirloctocogene samoparvovec (SPK-8011) | $4.26 \times 10^{-13}$ | ↓38.9 | $3.84 \times 10^{-12}$ | ↓4.31 | $1.66 \times 10^{-11}$ |

Using the predicted human GEF derived from preclinical data and targeted plasma FVIII activity level (12 IU/dL) or targeted FIX level (27.5 ng/mL for FIX-Pauda and 250 ng/mL for FIX), human effective starting doses were predicted using body weight-based direct conversion, AS-logW, and AS-W^-0.25^ approaches (Table 2 and Table S4). Among the 9 vectors, the FIP doses derived from AS-logW were 1.60 – 9.9 fold higher than that derived from AS-W^-0.25^. Except for vectors rAAV2-hAAT-FIX and SPK-9001, body weight-based direct conversion generated either comparable or lower predicted FIP doses than AS-W^-0.25^.

**Table 2.**
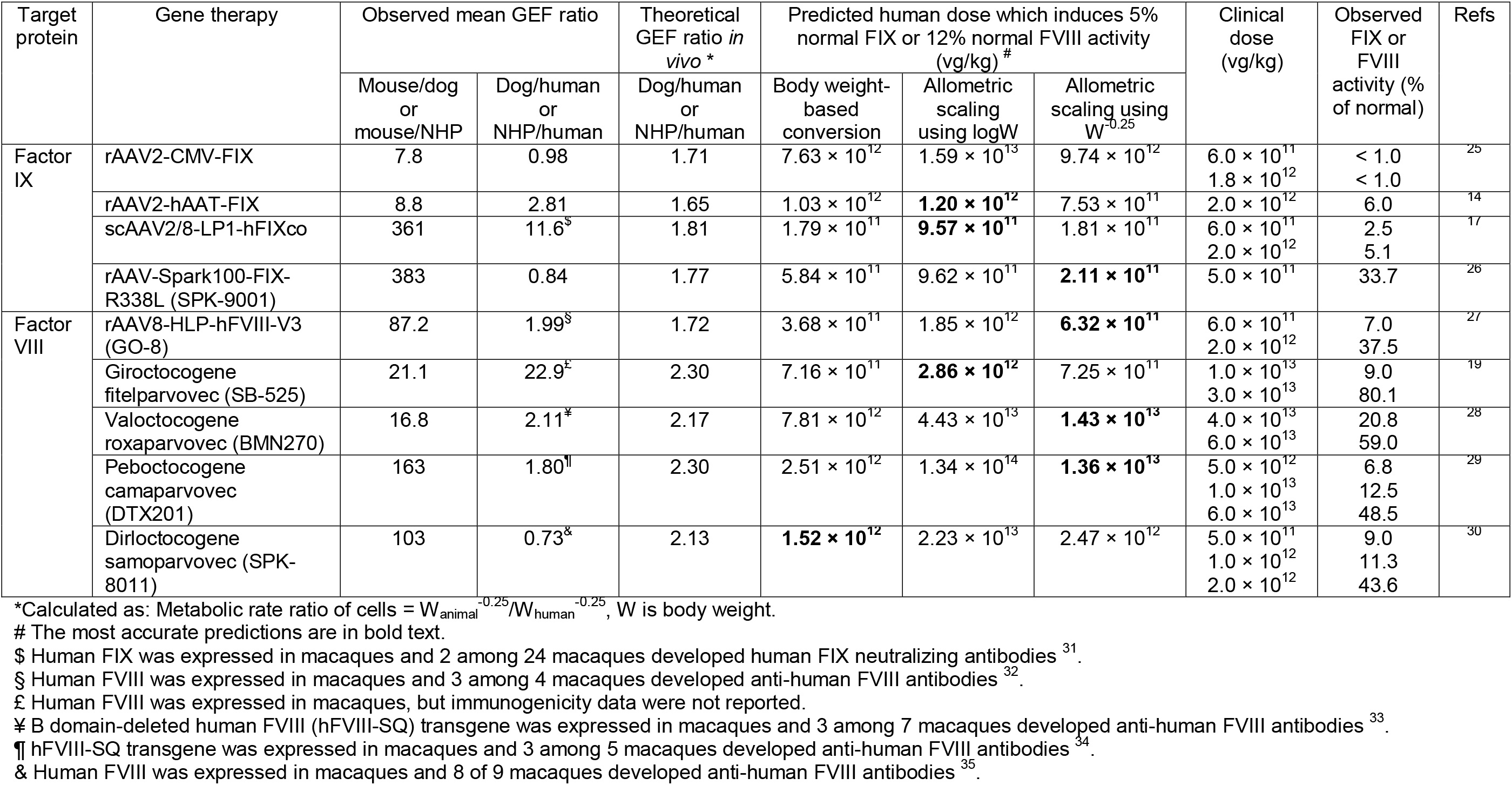
A comparison of two allometric scaling approaches and direct body weight-based conversion approach for human dose predictions

The FIP doses predicted by the three approaches were compared with clinical doses (Table 2). For vector rAAV2-CMV-FIX, even at the highest clinical dose 1.8×10^12^ vg/kg, the FIX level in patients was less than 1% of normal. The three predicted FIH doses were higher than 1.8×10^12^ vg/kg. It is not feasible to determine which predicted FIP dose is more likely to achieve the targeted FIX level (5% of normal). Among the other 8 vectors, AS-W^-0.25^ approach generated more accurate FIP dose predictions for 4 vectors (SPK-9001, GO-8, BMN270 and DTX201), AS-logW approach provided more accurate predictions for three vectors (rAAV2-hAAT-FIX, scAAV2/8-LP1-hFIXco and SB-525), and body weight-based direct conversion gave a more accurate prediction for SPK-8011.

The foundation of allometric scaling is the inverse relationship between metabolic rate of cells in vivo and W^0.25^ of mammalian species ^2^. Since protein and DNA/RNA synthesis is shown to be most sensitive to cellular energy supply ^16^, it appears reasonable to expect that the theoretical GEF is reversely correlated with the W^0.25^ of mammalian species. Based on the body weight of dogs, monkeys and hemophilia patients in each study, the theoretical intracellular metabolic rate ratio or GEF ratio in vivo between dog or monkey and patient was determined between 1.65 – 2.30 (Table 2). However, the observed dog/human or monkey/human GEF ratios for some vectors from corresponding theoretical GEF ratio. When the observed monkey/human or dog/human GEF ratio was much higher than the theoretical GEF ratio (i.e., rAAV2-hAAT-FIX, scAAV2/8-LP1-hFIXco, and SB-525), a high human dose was required. All the three approaches underestimated FIP doses but AS-logW approach provided more accurate predictions. When the observed monkey/human GEF ratio or dog/human GEF ratio was equal or less than 1 (SPK-9001 and SPK-8011), all the three approaches overestimated FIP doses but either body weight-based conversion or AS-W^-0.25^ provided more accurate predictions. Following the treatment of rAAV2-CMV-FIX vector, the plasma FIX levels in patients were less than 1% of normal, which likely included both baseline endogenous FIX and transgene product. Therefore, the observed human GEF (66.8 molecules/day/vg, Table 1) of rAAV2-CMV-FIX was likely overestimated and the observed dog/human GEF ratio of rAAV2-CMV-FIX (0.98) was likely underestimated. Although the observed dog/human GEF ratio of rAAV2-CMV-FIX was less than 1, we could not determine if the three approaches over- or under-estimated FIP dose. When the observed monkey/human GEF ratio or dog/human GEF ratio was similar with the theoretical ratio (GO-8, BMN270 and DTX201), not surprisingly, AS-W^-0.25^ provided more accurate predictions of FIP dose.

The performance of the three approaches was dependent on the observed monkey/human GEF ratio or dog/human GEF ratio. We explored if preclinical or in vitro data could be used to predict monkey/human GEF ratio or dog/human GEF ratio. Unfortunately, a relationship between mouse/dog or mouse/monkey GEF ratios and dog/human or monkey/human GEF ratios was not observed (Table 2). Among the three vectors with a high observed dog/human or monkey/human GEF ratio (i.e., rAAV2-hAAT-FIX, scAAV2/8-LP1-hFIXco, and SB-525), patients receiving high dose of rAAV2-hAAT-FIX ^14, 15^ or medium-to-high dose of scAAV2/8-LP1-hFIXco ^17, 18^ reported CD^8+^ T-cell responses against capsid, which could eliminate vector-transduced hepatocytes (Table S7). The preliminary efficacy and safety data of SB-525 was published as a conference abstract ^19^. Eight among 11 patients showed an increase in alanine aminotransferease (ALT) following the treatment of SB-525. It is unknown if T-cell responses to SB-525 were tested or not in the study. T-cell responses against capsid were not reported in the clinical studies of the other eight vectors. The high observed dog/human and monkey/human GEF ratios might be due to T-cell responses in hemophilia patients.

Based on the reverse allometric relationship between intracellular GEF and W^0.25^ of mammalian specie, the GEFs in dogs and monkeys were expected to be higher than that in humans. However, two bio-engineered AAV capsids Spark100 and Spark200 utilized in SPK-9001 and SPK-8011, respectively, have a higher transduction efficiency for human hepatocytes than natural AAV serotypes ^10, 20^. Spark100 and Spark200 capsids were identified by screening a human specific replication competent viral library composed of DNA shuffled AAV capsids with a humanized mouse liver model. In the humanized mouse liver model, AAV-Spark200 (AAV-LK03) transduced primary human hepatocytes 100-fold higher than AAV8 ^20^. The high transduction efficiency of SPK-9001 and SPK-8011 for human hepatocytes might contribute to their high GEF in humans and low dog/human or monkey/human GEF ratio.

## Discussion

In this study, we compared the performance of three approaches for human GEF and FIP dose predictions for nine AAV vectors. Both AS-logW and AS-W^-0.25^ analyses demonstrated a reverse correlation between GEF and logW or W^-0.25^, indicating that body weight is an important intrinsic factor which affects the expression efficiency of target proteins following gene therapy. In general, preclinical and clinical GEF values correlated well with W^-0.25^ of difference species. However, intense immune responses (i.e., T-cell responses) in patients and the high selectivity of bio-engineered AAV capsids to human hepatocytes might complicate the relationship between GEF and W^-0.25^. All the three factors, including body weight, immune responses, and human-specific transduction efficiency, should be considered when we predict FIP dose using preclinical data.

Mice, dogs and monkeys are commonly used as the animal models for AAV-mediated gene therapy ^21^. Our allometric scaling analysis revealed the importance of body weight in GEF of gene therapy. Compared to the three preclinical species with body weight < 15 kg, the GEF in large animals (i.e., swine and cattle) is expected to be more representative of that in humans. Furthermore, including GEF data of a large animal model can improve the quality of allometric scaling based human GEF predictions. Although the production of AAV vectors for a large animal study may increase the costs of preclinical development, an accurate prediction of FIP dose can reduce the costs of clinical development and increase the chances of success in dose-finding study.

Immune responses in animals and humans are a big challenge in developing AAV-mediated gene therapy. Specifically, three immunogenic targets in AAV-mediated gene therapy have been identified and discussed: transgene product (FIX, FVIII, FIX-Pauda, B-domain deleted recombinant FVIII), AAV capsid, and AAV capsid antigens expressed on the surface of transduced cells ^10^. Anti-transgene product antibodies (FIX or FVIII inhibitors) have not been reported in hemophilia AAV clinical trials ^10^ but anti-human FIX or FVIII antibodies were often detected in monkeys and dogs receiving AAV encoding a human transgene product. As shown in Table 2, the vectors encoding human FIX or FVIII (scAAV2/8-LP1-hFIXco, GO-8, BMN270, DTX201 and SPK8011) induced anti-human FIX or FVIII antibodies in dogs or monkeys. The formation of human FIX or FVIII inhibitors can decrease the GEF in dogs and monkeys and reduce dog/human or monkey/human GEF ratio, which probably partially contributed to the low monkey/human GEF ratios of DTX201 and SPK-8011. To avoid the negative impact of FIX or FVIII inhibitors, preclinical FIX or FVIII data collected prior to the detection of inhibitors should be used to calculate the GEF of human FIX or FVIII in animals.

Anti-AAV neutralizing antibodies and AAV capsid specific T cells exist in many people because of prior exposure to the common wide-type virus ^10, 22^. AAV capsid antibodies may preclude transduction of AAV vectors and T-cell response may eliminate transduced human hepatocytes ^10^. Dose selection for patients with preexisting immunity prior to gene therapy is quite a dilemma. Preexisting immunity especially T-cell response can reduce the levels of transgene product to sub-therapeutic range and a high FIP dose of AAV vector is needed for patients with preexisting immunity. On the other hand, vector immunogenicity is dose-dependent and a high dose of AAV vector further increases the risk of intense immune responses in patients with preexisting immunity. Ideally, patients with preexisting immunity should be excluded from a first-in-human study of an intravenously administered liver-targeted gene therapy. When an effective and safe AAV dose has been determined in AAV naïve patients, the AAV dose can be further adjusted for patients with preexisting immunity based on individual’s levels of anti-AAV capsid antibodies and AAV capsid specific T cells. If it is not feasible to exclude patients with preexisting immunity in a first-in-human study (i.e., it is challenging to enroll AAV naïve patients due to a high prevalence of anti-AAV preexisting immunity), our analysis suggests that AS-logW approach is more likely to provide an effective FIP dose for patients who are expected to show intense immune responses compared to AS-W^-0.25^ and body weight-based direct conversion. The FIP dose can be predicted using both AS-logW and AS-W^-0.25^. A final dose between the two predicted values can be selected based on the titers of preexisting AAV neutralizing antibodies and the presence or absence of preexisting T cell responses.

Both rAAV2-hAAT-FIX and scAAV2/8-LP1-hFIXco triggered AAV capsid specific T-cell responses in patients receiving a medium or high dose of vector, which probably caused a substantial underprediction of FIP dose using all three approaches. The presence of immune response especially AAV capsid specific T-cell responses complicated FIP dose prediction. It will benefit FIP dose prediction if T-cell response risk of AAV vectors can be predicted using in vitro assays (i.e., enzyme-linked immunospot assay) ^23^ and humanized animal models prior to clinical study. More research in this field is warranted.

Different from wide-type AAV vectors, novel bioengineered AAV capsids often exhibit a similar or even higher transduction efficiency in human hepatocytes compared to that in preclinical species, which may cause an overprediction of FIP dose using traditional allometric scaling or body weight-based dose conversion approaches. The AAV transduction efficiency in animal and human hepatocytes can be assessed using a humanized mouse liver model ^20^. For these vectors with a high transduction efficiency in human hepatocytes, it appears appropriate to use body weight-based dose conversion for FIP dose prediction.

Another concern of AAV-mediated hemophilia gene therapy is the potential supra-physiologic levels of transgene product. Although the upper safety margin of FIX and FVIII in hemophilia patients receiving gene therapy is still unclear, FIX or FVIII levels above 150% of normal have been associated with thrombosis in normal subjects ^24^. If we assume 150% of normal FIX or FVIII level is the upper safety margin, the FIP doses for SPK-9001 and SPK-8011 predicted by the AS-W^-0.25^ approach are unlikely to generate supra-physiologic levels of transgene product even though the doses were overpredicted.

Based on the findings from this study, a FIP dose prediction decision tree is proposed for AAV-mediated hemophilia gene therapy (Figure 3). The decision tree tentatively proposes criteria for prediction approach selection. A successful prediction of an effective and safe FIP dose replies on removing of vectors with a high immunogenicity risk in humans at preclinical stage, eliminating the negative impact of immunogenicity on transgene product in preclinical species, excluding patients with preexisting immunity to AAV, and identifying the differences in vector transduction efficiency between dogs/monkeys and humans using humanize animal models.

**Figure 3.**
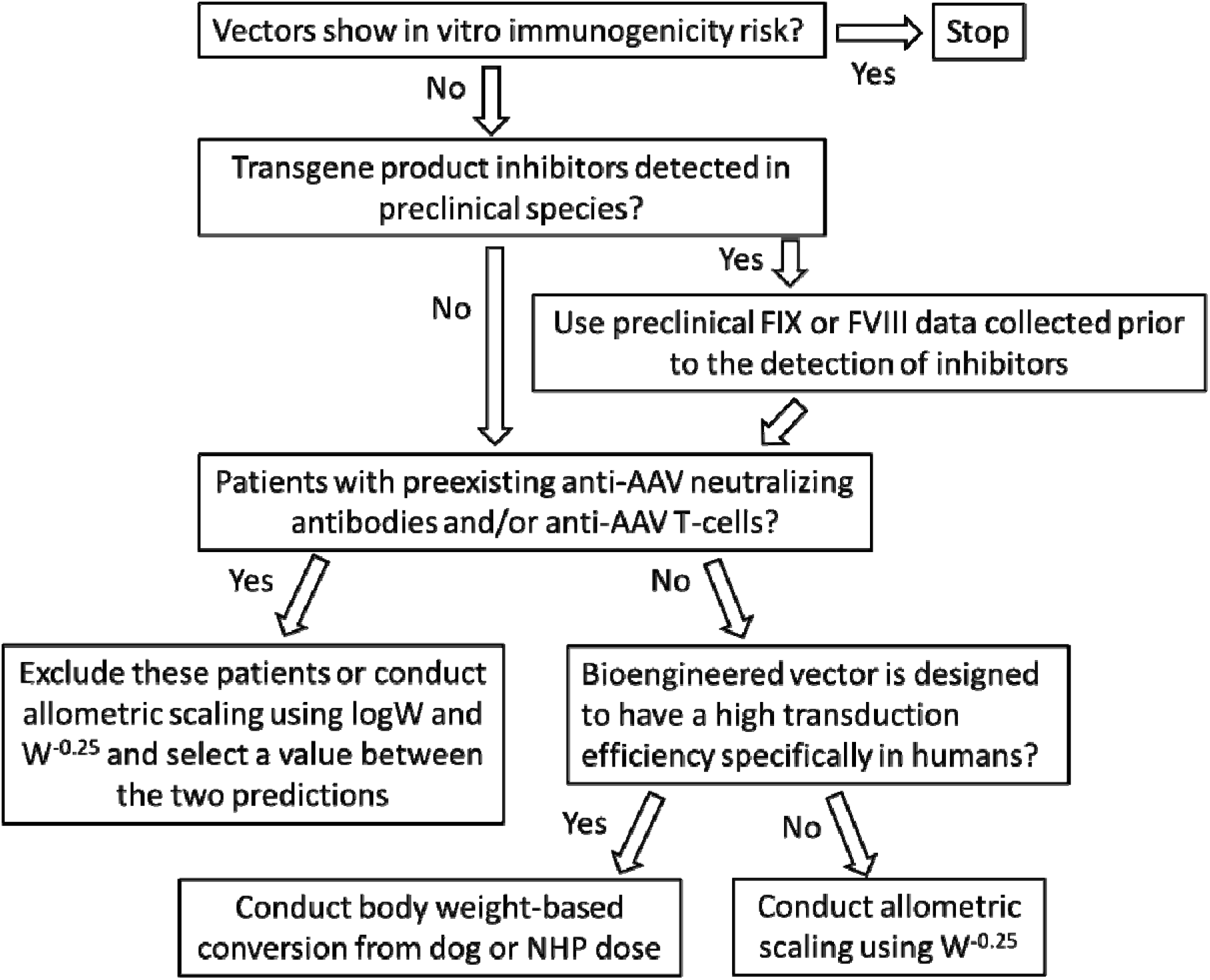
First-in-patient dose prediction decision tree for adeno-associated virus-mediated hemophilia gene therapy.

Finally, the limited public access to preclinical efficacy and safety data of AAV vectors is another obstacle to FIP dose prediction. Although the clinical results of most AAV-mediated gene therapies have been published, only a small fraction of AAV vectors have preclinical efficacy and safety results publicly available. People will be more confident about the predict FIP dose if the prediction approaches are validated with the data of more AAV vectors. A collaboration organization such as an IQ consortium may facilitate information exchange among researchers and stakeholders.

## Conclusions

In this study, the author compared the performance of two allometric scaling approaches (AS-logW and AS-W^-0.25^) and body weight-based dose conversion approach for FIP dose prediction for AAV-mediated hemophilia gene therapy. The prediction performance of the three approaches was assessed using preclinical and clinical efficacy data of nine AAV vectors. In general, body weight-based direct conversion approach was more likely to underestimate FIP dose but worked better for one bioengineered vector with a high human hepatocyte transduction efficiency. In contrast, AS-logW was likely to overestimate FIP dose but worked better than the other two approaches when vector capsid-specific T-cell responses were detected in patients. The third approach, AS-W^-0.25^, was appropriate for FIP dose prediction in the absence of T-cell responses to AAV vectors or a dramatic difference in vector transduction efficiency between animals and humans. A decision tree was tentatively proposed to facilitate approach selection for FIP dose prediction in AAV-mediated hemophilia gene therapy. Three factors including body weight of preclinical species and patients, immune responses, and human-specific transduction efficiency should be considered when we predict FIP dose of AAV-mediate hemophilia gene therapy using preclinical data.

## Notes

### Competing Interest Statement

The authors have declared no competing interest.

