## Supplementary materials for "First-in-patient dose prediction for adeno-associated virus-mediated hemophilia gene therapy using allometric scaling"

Peng Zou*

Quantitative Clinical Pharmacology, Daiichi Sankyo, Inc., 211 Mt. Airy Road, Basking Ridge, NJ 07920

*Corresponding author

Peng Zou, Ph.D.

Quantitative Clinical Pharmacology

Daiichi Sankyo, Inc.

211 Mount Airy Road

Basking Ridge, NJ 07920

**Figure S1. Allometric scaling of gene efficiency factor (GEF) for (A and B) rAAV2-CMV-FIX, (C and D) rAAV2-hAAT-FIX, (E and F) scAAV2/8-LP1-hFIXco, and (G and H) rAAV-Spark100-FIX-R338L using preclinical data from two species.** The equations were used to predict human GEF. LogW was used as the scaling variable in A, C and E and W^-0.25^ was used as the scaling variable in B, D and F. W is body weight in kg and the unit of GEF is molecules/day/viral genome. Circle, triangle, and square symbols represent mouse, dog, and macaque, respectively.

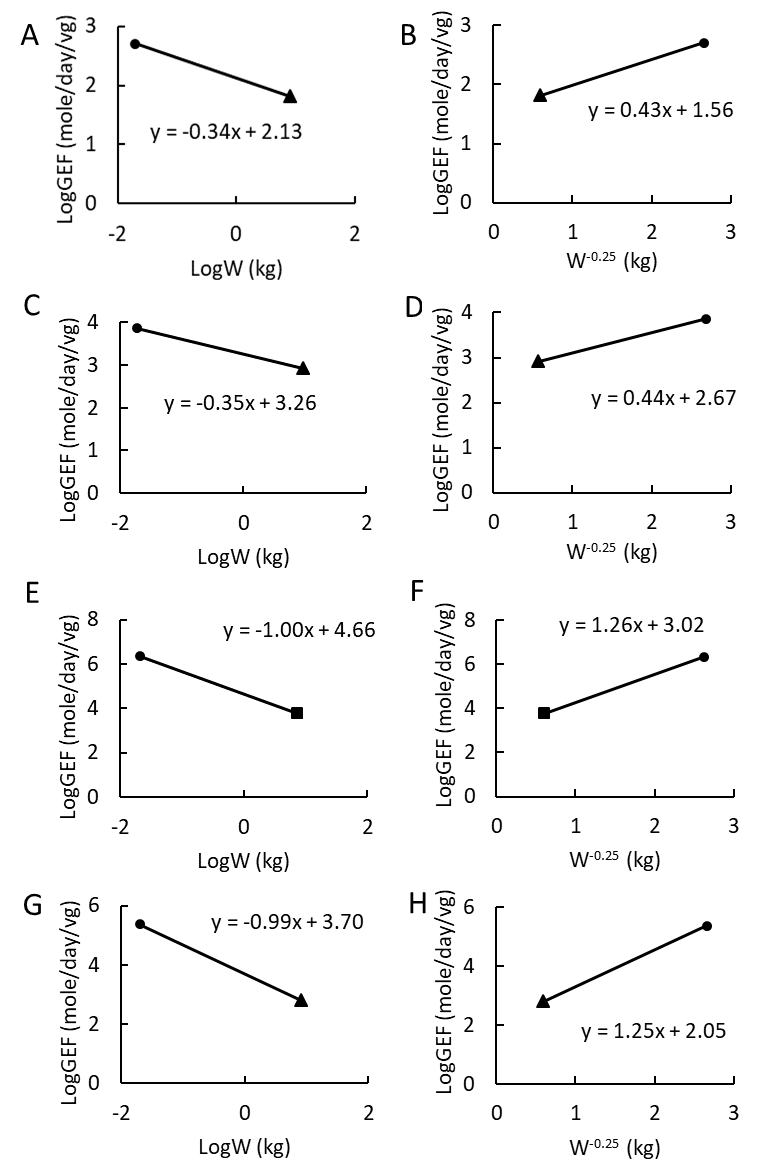

**Figure S2. Allometric scaling of gene efficiency factor (GEF) for (A and B) GO-8, (C and D) SB-525, (E and F) BMN270, (G and H) DTX201, and (I and J) SPK-8011** **using preclinical data from two species.** The equations were used to predict human GEF. LogW was used as the scaling variable in A, C, E, G, and I and W^-0.25^ was used as the scaling variable in B, D, F, H, and J. W is body weight in kg and the unit of GEF is IU/day/viral genome. Circle and square symbols represent mouse, macaque, and human, respectively.

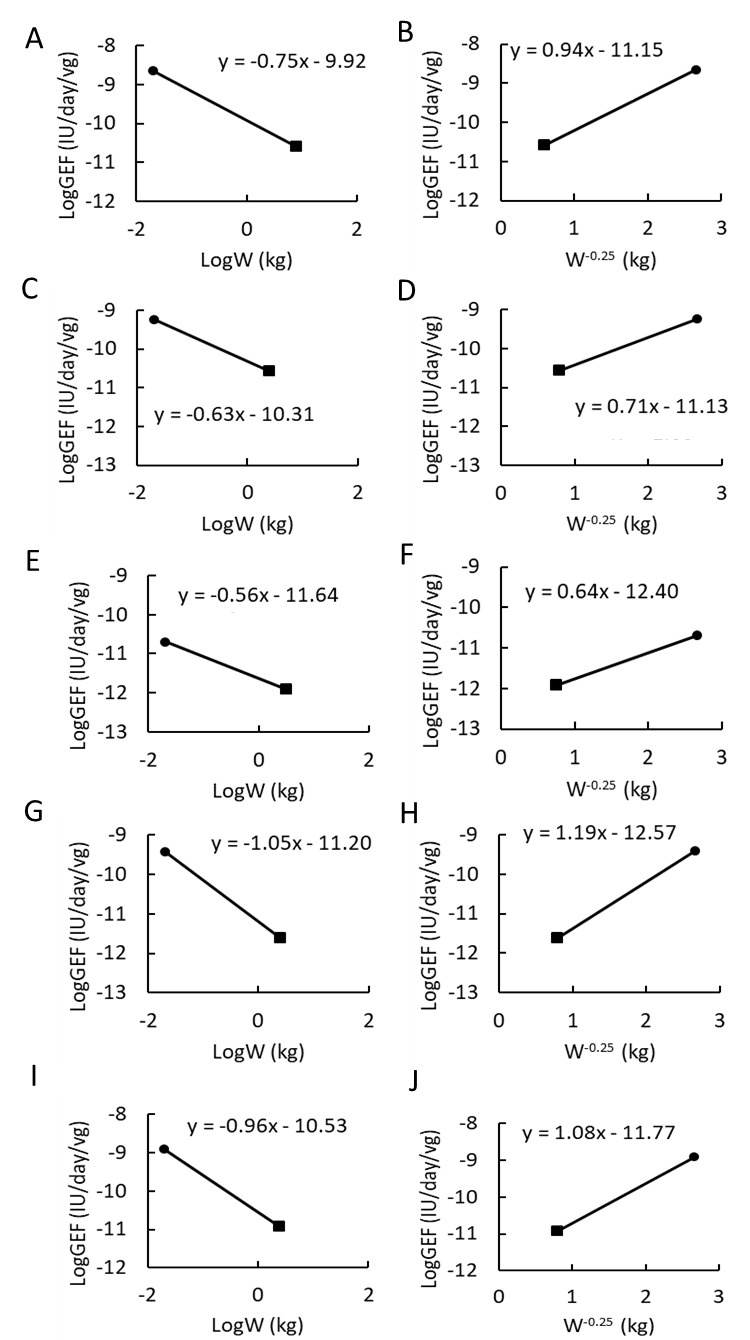

**Table S1. Plasma FVIII activity and gene efficiency factor values used for allometric analysis for FVIII vectors.**

| Vector | Species | Dose (vg/kg) | Body weight (kg) | Plasma FVIII activity (IU/dL or % of normal) | GEF (IU/day/vg) | Refs |
| --- | --- | --- | --- | --- | --- | --- |
| rAAV8-HLP-hFVIII-V3 (GO-8) | Mice | 2.00E+12 | 0.02 | 732 | 3.32E-09 | ^1^ |
|  |  | 2.00E+13 | 0.02 | 2578 | 1.17E-09 |  |
|  | Monkeys (Rhesus) | 2.00E+12 | 8 | 43 | 2.69E-11 | ^1^ |
|  |  | 7.00E+12 | 8 | 138 | 2.47E-11 |  |
|  | Humans | 6.00E+11 | 70 | 7 | 9.24E-12 | ^2^ |
|  |  | 2.00E+12 | 70 | 6 | 2.38E-12 |  |
|  |  | 2.00E+12 | 70 | 69 | 2.73E-11 |  |
| SB-525 | Mice | 7.20E+12 | 0.02 | 458.1 | 5.78E-10 | ^3^ |
|  | Monkeys (Cynomolgus) | 2.00E+11 | 2.5 | 3.9 | 2.44E-11 | ^3^ |
|  |  | 6.00E+11 | 2.5 | 6.4 | 1.33E-11 |  |
|  |  | 9.00E+11 | 2.5 | 11.7 | 1.63E-11 |  |
|  |  | 2.00E+12 | 2.5 | 56.4 | 3.53E-11 |  |
|  |  | 6.00E+12 | 2.5 | 227.9 | 4.75E-11 |  |
|  | Humans | 2.00E+12 | 70 | 3.1 | 1.23E-12 | ^4^ |
|  |  | 1.00E+13 | 70 | 5 | 3.96E-13 |  |
|  |  | 1.00E+13 | 70 | 13 | 1.03E-12 |  |
|  |  | 3.00E+13 | 70 | 80.1 | 2.11E-12 |  |
| BMN270 | Mice | 2.00E+12 | 0.02 | 4.69 | 2.13E-11 | ^5^ |
|  |  | 2.00E+13 | 0.02 | 23.5 | 1.07E-11 |  |
|  |  | 2.00E+14 | 0.02 | 287 | 1.30E-11 |  |
|  |  | 6.00E+12 | 0.02 | 4.9 | 7.41E-12 |  |
|  |  | 2.00E+13 | 0.02 | 53 | 2.41E-11 |  |
|  |  | 6.00E+13 | 0.02 | 299 | 4.52E-11 |  |
|  | Monkeys (Cynomolgus) | 1.00E+13 | 3.15 | 22.8 | 2.85E-12 | ^5, 6^ |
|  |  | 1.00E+13 | 3.15 | 4.8 | 6.00E-13 |  |
|  |  | 3.60E+13 | 3.15 | 41.3 | 1.43E-12 |  |
|  |  | 3.6E+13 | 3.15 | 45.6 | 1.58E-12 |  |
|  |  | 6.00E+13 | 3.15 | 32.9 | 6.86E-13 |  |
|  |  | 6.00E+13 | 3.15 | 38.2 | 7.96E-13 |  |
|  |  | 6.00E+13 | 3.15 | 24.3 | 5.06E-13 |  |
|  | Humans | 2.00E+13 | 103 | 2 | 7.92E-14 | ^7, 8^ |
|  |  | 4.00E+13 | 70 | 18 | 3.56E-13 |  |
|  |  | 4.00E+13 | 70 | 40 | 7.92E-13 |  |
|  |  | 4.00E+13 | 70 | 24 | 4.75E-13 |  |
|  |  | 4.00E+13 | 70 | 14 | 2.77E-13 |  |
|  |  | 4.00E+13 | 70 | 26 | 5.15E-13 |  |
|  |  | 4.00E+13 | 70 | 3 | 5.94E-14 |  |
|  |  | 6.00E+13 | 70 | 62 | 8.18E-13 |  |
|  |  | 6.00E+13 | 70 | 50 | 6.60E-13 |  |
|  |  | 6.00E+13 | 70 | 95 | 1.25E-12 |  |
|  |  | 6.00E+13 | 70 | 11 | 1.45E-13 |  |
|  |  | 6.00E+13 | 70 | 88 | 1.16E-12 |  |
|  |  | 6.00E+13 | 70 | 52 | 6.86E-13 |  |
|  |  | 6.00E+13 | 70 | 55 | 7.26E-13 |  |
| DTX201 | Mice | 3.00E+11 | 0.02 | 17.3 | 5.24E-10 | ^9^ |
|  |  | 1.00E+12 | 0.02 | 60 | 5.45E-10 |  |
|  |  | 3.00E+12 | 0.02 | 110 | 3.33E-10 |  |
|  |  | 1.00E+13 | 0.02 | 168.4 | 1.53E-10 |  |
|  | Monkeys (Cynomolgus) | 1.20E+13 | 2.5 | 37.1 | 3.87E-12 | ^10^ |
|  |  | 1.20E+13 | 2.5 | 10.5 | 1.09E-12 |  |
|  |  | 1.20E+13 | 2.5 | 11 | 1.15E-12 |  |
|  |  | 1.20E+13 | 2.5 | 27 | 2.81E-12 |  |
|  |  | 1.20E+13 | 2.5 | 29 | 3.02E-12 |  |
|  | Humans | 5.00E+12 | 70 | 10 | 1.58E-12 | ^11^ |
|  |  | 5.00E+12 | 70 | 3.5 | 5.54E-13 |  |
|  |  | 1.00E+13 | 70 | 20 | 1.58E-12 |  |
|  |  | 1.00E+13 | 70 | 5 | 3.96E-13 |  |
|  |  | 2.00E+13 | 70 | 73 | 2.89E-12 |  |
|  |  | 2.00E+13 | 70 | 24 | 9.50E-13 |  |
| SPK-8011 | Mice | 4.00E+12 | 0.02 | 550 | 1.25E-09 | ^12^ |
|  | Monkeys (Cynomolgus) | 2.00E+12 | 4 | 22.3 | 1.39E-11 | ^12^ |
|  |  | 6.00E+12 | 4 | 62 | 1.29E-11 |  |
|  |  | 2.00E+13 | 4 | 153 | 9.57E-12 |  |
|  | Humans | 5.00E+11 | 68 | 10 | 1.58E-11 | ^13^ |
|  |  | 5.00E+11 | 89 | 8 | 1.27E-11 |  |
|  |  | 1.00E+12 | 89 | 5 | 3.96E-12 |  |
|  |  | 1.00E+12 | 82 | 15 | 1.19E-11 |  |
|  |  | 1.00E+12 | 89 | 14 | 1.11E-11 |  |
|  |  | 2.00E+12 | 60 | 16 | 6.34E-12 |  |
|  |  | 2.00E+12 | 79 | 37 | 1.47E-11 |  |
|  |  | 2.00E+12 | 93 | 20 | 7.92E-12 |  |
|  |  | 2.00E+12 | 72 | 8 | 3.17E-12 |  |
|  |  | 2.00E+12 | 78 | 15 | 5.94E-12 |  |
|  |  | 2.00E+12 | 78 | 46 | 1.82E-11 |  |
|  |  | 2.00E+12 | 83 | 16 | 6.34E-12 |  |
|  |  | 2.00E+12 | 69 | 117 | 4.63E-11 |  |
|  |  | 2.00E+12 | 121 | 117 | 4.63E-11 |  |
|  |  | 1.50E+12 | 60 | 29 | 1.53E-11 |  |
|  |  | 1.50E+12 | 115 | 29 | 1.53E-11 |  |
|  |  | 1.50E+12 | 60 | 38 | 2.01E-11 |  |
|  |  | 1.50E+12 | 95 | 70 | 3.70E-11 |  |

**Table S2. Plasma FIX activity and gene efficiency factor values used for allometric analysis for FIX vectors.**

| Vector | Species | Dose (vg/kg) | Body weight (kg) | Plasma FIX (ng/mL) | GEF (molecules/day/vg) | Refs |
| --- | --- | --- | --- | --- | --- | --- |
| rAAV2-CMV-FIX | Mice | 6.25E+11 | 0.016 | 98.3 | 1440 | ^14^ |
|  |  | 1.25E+13 | 0.016 | 286 | 209 |  |
|  |  | 4.44E+12 | 0.0225 | 170 | 350 |  |
|  |  | 4.44E+12 | 0.0225 | 140 | 288 |  |
|  |  | 4.44E+12 | 0.0225 | 135 | 278 |  |
|  |  | 2.00E+11 | 0.02 | 19 | 868 |  |
|  |  | 1.00E+12 | 0.02 | 20 | 183 |  |
|  |  | 4.00E+12 | 0.02 | 53 | 121 |  |
|  | Dogs | 1.30E+11 | 5.7 | 2.6 | 52.3 | ^14^ |
|  |  | 1.10E+12 | 9.1 | 12 | 28.5 |  |
|  |  | 3.40E+12 | 20 | 17 | 13.1 |  |
|  |  | 3.00E+12 | 13.6 | 21 | 18.3 |  |
|  |  | 8.50E+12 | 4.9 | 69 | 16.6 |  |
|  |  | 8.50E+12 | 4.9 | 39 |  |  |
|  |  | 5.60E+12 | 4.5 | 40 | 18.7 |  |
|  |  | 1.00E+12 | 8.4 | 34 | 89.0 |  |
|  |  | 3.00E+12 | 9 | 10 | 8.72 |  |
|  |  | 3.00E+12 | 9 | 11 | 9.59 |  |
|  |  | 2.20E+12 | 5.5 | 115 | 138 |  |
|  | Humans | 2.00E+11 | 70 | 10 | 118 | ^14^ |
|  |  | 2.00E+11 | 70 | 13.8 |  |  |
|  |  | 6.00E+11 | 70 | 16.7 | 63.6 |  |
|  |  | 6.00E+11 | 70 | 30 |  |  |
|  |  | 6.00E+11 | 70 | 10.8 |  |  |
|  |  | 1.80E+12 | 70 | 20 | 18.4 |  |
|  |  | 1.80E+12 | 70 | 13.3 |  |  |
| rAAV2-hAAT-FIX | Mice | 1.67E+13 | 0.018 | 8410 | 4610 | ^14^ |
|  |  | 4.00E+12 | 0.02 | 9630 | 22000 |  |
|  |  | 1.04E+12 | 0.0193 | 1060 | 9360 |  |
|  |  | 1.04E+13 | 0.0193 | 10900 | 9630 |  |
|  |  | 1.04E+14 | 0.0193 | 58500 | 5160 |  |
|  |  | 5.18E+12 | 0.0193 | 2750 | 4850 |  |
|  |  | 1.85E+11 | 0.02 | 4.07 | 201 |  |
|  |  | 5.50E+11 | 0.02 | 103 | 1710 |  |
|  |  | 1.65E+12 | 0.02 | 897 | 4970 |  |
|  |  | 5.50E+12 | 0.02 | 2820 | 5160 |  |
|  |  | 1.50E+13 | 0.02 | 9550 | 5820 |  |
|  |  | 5.00E+12 | 0.02 | 350 | 639 |  |
|  |  | 5.00E+12 | 0.02 | 87.1 | 159 |  |
|  |  | 5.00E+12 | 0.02 | 60.9 | 111 |  |
|  | Dogs | 1.20E+12 | 10.2 | 590 | 1260 | ^14^ |
|  |  | 1.60E+12 | 6 | 220 | 356 |  |
|  |  | 8.00E+11 | 12.3 | 262 | 878 |  |
|  | Humans | 2.00E+12 | 70 | 444 | 442 | ^15^ |
|  |  | 2.00E+12 | 70 | 150 | 149* |  |
| scAAV2/8-LP1-hFIXco | Mice | 5.00E+10 | 0.02 | 563 | 103000 | ^14^ |
|  |  | 1E+11 | 0.02 | 3250 | 297000 |  |
|  |  | 1.25E+12 | 0.02 | 36600 | 268000 |  |
|  |  | 5E+12 | 0.02 | 155000 | 284000 |  |
|  |  | 2.00E+11 | 0.0227 | 30400 | 1390000 |  |
|  |  | 4.00E+10 | 0.0227 | 17700 | 4040000 |  |
|  |  | 4.00E+09 | 0.0227 | 1610 | 3680000 |  |
|  |  | 4.00E+08 | 0.0227 | 322 | 7350000 |  |
|  | Monkeys (Cynomolgus) | 1.00E+12 | 4.9 | 1400 | 3100 | ^14^ |
|  |  | 1.00E+12 | 5.7 | 800 | 1770 |  |
|  |  | 1.00E+12 | 4.3 | 700 | 1550 |  |
|  |  | 2.00E+12 | 7.4 | 16100 | 17900 |  |
|  |  | 2.00E+12 | 8.5 | 3000 | 3320 |  |
|  |  | 2.00E+12 | 11.3 | 11000 | 12200 |  |
|  |  | 2.00E+11 | 11.7 | 523 | 5780 |  |
|  |  | 2.00E+11 | 15.1 | 1030 | 14400 |  |
|  |  | 2.00E+11 | 15.7 | 1490 | 16400 |  |
|  |  | 6.00E+10 | 6.7 | 161 | 5940 |  |
|  |  | 6.00E+10 | 7.3 | 213 | 7850 |  |
|  |  | 6.00E+10 | 6.3 | 235 | 8660 |  |
|  | Humans | 2.00E+11 | 80.7 | 109 | 889 | ^14^ |
|  |  | 2.00E+11 | 80.7 | 70 |  |  |
|  |  | 6.00E+11 | 80.7 | 143 | 417 |  |
|  |  | 6.00E+11 | 80.7 | 109 |  |  |
|  |  | 2.00E+12 | 80.7 | 178 | 254 |  |
|  |  | 2.00E+12 | 80.7 | 361 |  |  |
|  |  | 2.00E+12 | 80.7 | 250 |  |  |
|  |  | 2.00E+12 | 80.7 | 334 |  |  |
|  |  | 2.00E+12 | 80.7 | 262 |  |  |
|  |  | 2.00E+12 | 80.7 | 145 |  |  |
| rAAV-Spark100-FIX-R338L (SPK-9001) | Mice | 1.00E+10 | 0.02 | 53.8 | 270180 | ^16^ |
|  |  | 4.00E+10 | 0.02 | 173.5 | 217826 |  |
|  |  | 1.00E+11 | 0.02 | 424.1 | 212980 |  |
|  |  | 4.00E+11 | 0.02 | 1978.5 | 248398 |  |
|  | Dogs | 1.00E+12 | 8.6 | 39.9 | 573 | ^16^ |
|  |  | 3.00E+12 | 10.2 | 241.2 | 1156 |  |
|  |  | 3.00E+12 | 5.8 | 26.8 | 128 |  |
|  | Humans | 5.00E+11 | 81.8 | 295.6 | 656 | ^17^ |
|  |  | 5.00E+11 | 55.6 | 953.3 | 788 |  |
|  |  | 5.00E+11 | 97.9 | 2330.2 | 547 |  |
|  |  | 5.00E+11 | 101.3 | 10870.9 | 897 |  |
|  |  | 5.00E+11 | 87 | 219.2 | 788 |  |
|  |  | 5.00E+11 | 70.9 | 1325.3 | 394 |  |
|  |  | 5.00E+11 | 87 | 147.3 | 306 |  |
|  |  | 5.00E+11 | 68.2 | 164.8 | 591 |  |
|  |  | 5.00E+11 | 82.8 | 197.8 | 1772 |  |
|  |  | 5.00E+11 | 68.1 | 137.4 | 634 |  |
| AMT-060 | Monkeys (Cynomolgus) | 5.00E+12 | 2.5 | 245 | 103.8 | ^18^ |
|  | Humans | 5.00E+12 | 84.5 | 220 | 87.6 | ^19^ |
|  |  | 2.00E+13 | 84 | 345 | 34.3 |  |
| rAAV5- FIX-R338L (AMT-061) | Monkeys (Cynomolgus) | 5.00E+11 | 2.5 | 100 | 442 | ^18^ |
|  |  | 5.00E+12 | 2.5 | 243 | 243 |  |
|  |  | 2.50E+13 | 2.5 | 1500 | 133 |  |
|  |  | 9.00E+13 | 2.5 | 3000 | 73.7 |  |
|  | Humans | 2.00E+13 | 89 | 392 | 39.1 | ^20, 21^ |
|  |  | 2.00E+13 | 81 | 255 | 25.4 |  |
|  |  | 2.00E+13 | 82 | 438 | 43.6 |  |
|  |  | 2.00E+13 | 70 | 318 | 31.6 |  |

*Subject F with plasma FIX level of 150 ng/mL (3% of normal level) was included in the calculation ^15^.

**Table S3. Clearance values of recombinant factor IX and factor VIII used for gene efficiency factor calculations.**

| Species | CL_FIX_ (mL/h/kg) | CL_FVIII_ (mL/h/kg) |
| --- | --- | --- |
| Mice | 34.8 ± 3.1 ^14^ | 37.83 ± 36.1 ^22-24^ |
| Dogs | 9.96 ± 0.55 ^14^ | N.A. |
| Monkeys | 8.42 ± 2.00 ^14^ | 5.21 ^24^ |
| Humans | 7.58 ± 1.84 ^14^ | 3.30 ± 0.28 ^22, 25^ |

| Vectors and refs | Species | Weight (kg) | Observed mean GEF (molecule/day/vg for FIX and IU/day/vg for FVIII) | Predicted human GEF (ng/day/vg for FIX and IU/day/vg for FVIII) | | Human CL | Target plasma FIX or FVIII level | Predicted human dose (vg/kg) | |
| --- | --- | --- | --- | --- | --- | --- | --- | --- | --- |
|  |  |  |  | logW | W^-0.25^ |  |  | logW | W^-0.25^ |
| rAAV2-CMV-FIX ^14^ | Mouse | 0.02 | 507 | N.A. | N.A. | 7.58 mL/h/kg | 250 ng/mL or 5% of normal plasma level | N.A. | N.A. |
|  | Dog | 8.1 | 65.3 | N.A. | N.A. |  |  | N.A. | N.A. |
|  | Human | 70 | 66.8 | 2.857E-9 | 4.670E-9 |  |  | 1.59E+13 | 9.74E+12 |
| rAAV2-hAAT-FIX ^14^ | Mouse | 0.019 | 7290 | N.A. | N.A. |  |  | N.A. | N.A. |
|  | Dog | 9.5 | 831 | N.A. | N.A. |  |  | N.A. | N.A. |
|  | Human | 70 | 296* | 3.780E-8 | 6.041E-8 |  |  | 1.20E+12 | 7.53E+11 |
| scAAV2/8-LP1-hFIXco ^14^ | Mouse | 0.021 | 2180000 | N.A. | N.A. |  |  | N.A. | N.A. |
|  | Monkey (Rhesus) | 7.5 | 6040 | N.A. | N.A. |  |  | N.A. | N.A. |
|  | Human | 81 | 520 | 4.752E-8 | 2.508E-7 |  |  | 9.57E+11 | 1.81E+11 |
| rAAV-Spark100-FIX-R338L (SPK-9001) | Mouse | 0.02 | 237346 | N.A. | N.A. |  | 27.5 ng/mL or 5% of normal plasma level | N.A. | N.A. |
|  | Dog | 8.2 | 619 | N.A. | N.A. |  |  | N.A. | N.A. |
|  | Human | 80 | 737 | 5.404E-8 | 2.463E-7 |  |  | 9.62E+11 | 2.11E+11 |
| GO-8 (FVIII) ^1, 2^ | Mouse | 0.02 | 2.24665E-09 | N.A. | N.A. | 3.30 mL/h/kg | 12 IU/dL or 12% of normal plasma level | N.A. | N.A. |
|  | Monkey (Rhesus) | 8 | 2.57672E-11 | N.A. | N.A. |  |  | N.A. | N.A. |
|  | Human | 70 | 1.298E-11 | 5.112E-12 | 1.503E-11 |  |  | 1.86E+12 | 6.32E+11 |
| SB-525 (FVIII) ^3, 4^ | Mouse | 0.02 | 5.77664E-10 | N.A. | N.A. |  |  | N.A. | N.A. |
|  | Monkey (Cynomolgus) | 2.5 | 2.73462E-11 | N.A. | N.A. |  |  | N.A. | N.A. |
|  | Human | 70 | 1.19196E-12 | 3.329E-12 | 1.311E-11 |  |  | 2.86E+12 | 7.25E+11 |
| BMN270 (FVIII) ^5, 6, 7, 8^ | Mouse | 0.02 | 2.02844E-11 | N.A. | N.A. |  |  | N.A. | N.A. |
|  | Monkey (Cynomolgus) | 3.15 | 1.20822E-12 | N.A. | N.A. |  |  | N.A. | N.A. |
|  | Human | 70 | 5.71843E-13 | 2.145E-13 | 6.636E-13 |  |  | 4.43E+13 | 1.43E+13 |
| DTX201 (FVIII) ^9-11^ | Mouse | 0.02 | 3.88529E-10 | N.A. | N.A. |  |  | N.A. | N.A. |
|  | Monkey (Cynomolgus) | 2.5 | 2.38826E-12 | N.A. | N.A. |  |  | N.A. | N.A. |
|  | Human | 70 | 1.3266E-12 | 7.115E-14 | 7.001E-13 |  |  | 1.34E+14 | 1.36E+13 |
| SPK-8011 (FVIII) ^12, 13^ | Mouse | 0.02 | 1.24839E-09 | N.A. | N.A. |  |  | N.A. | N.A. |
|  | Monkey (Cynomolgus) | 2.5 | 1.21428E-11 | N.A. | N.A. |  |  | N.A. | N.A. |
|  | Human | 82 | 1.65733E-11 | 4.263E-13 | 3.844E-12 |  |  | 2.23E+13 | 2.47E+12 |

**Table S4. Mean observed preclinical and clinical FIX or FVIII gene efficiency factor (GEF) values, predicted human GEF, and predicted human doses for targeted protein levels.**

*Subject F with plasma FIX level of 150 ng/mL (3% of normal level) was included in the calculation ^15^.

**Table S5. LogW-dependent allometric equations and human predictions for gene efficiency factor (GEF)**

| Vector | Allometric Equation using log_10_W | | Human GEF (molecule/day/vg for FIX) (IU/day/vg for FVIII) | |
| --- | --- | --- | --- | --- |
|  | Three species | Two species | Observed | Predicted |
| rAAV2-CMV-FIX | logGEF = -0.268 * logW  + 2.209 (r^2^ = 0.9304) | logGEF = -0.341 * logW + 2.125 | 66.8 | 31.3 |
| rAAV2-hAAT-FIX | logGEF = -0.380 * logW  + 3.224 (r^2^ = 0.993) | logGEF = -0.349 * logW + 3.261 | 296 | 414 |
| scAAV2/8-LP1-hFIXco | logGEF = -1.009 * logW  + 4.650 (r^2^ = 1.00) | logGEF = -1.002 * logW + 4.658 | 520 | 557 |
| rAAV-Spark100-FIX-R338L (SPK-9001) | logGEF = -0.756 * logW  + 3.960 (r^2^ = 0.916) | logGEF = -0.989 * logW + 3.695 | 737 | 65.0 |
| GO-8 | logGEF = -0.656 * logW  – 9.812 (r^2^ = 0.981) | logGEF = -0.746 * logW – 9.915 | 1.30 × 10^-11^ | 5.11 × 10^-12^ |
| SB-525 | logGEF = -0.749* logW  – 10.439 (r^2^ = 0.987) | logGEF = -0.632 * logW – 10.312 | 1.19 × 10^-12^ | 3.33 × 10^-12^ |
| BMN270 | logGEF = -0.449 * logW  – 11.521 (r^2^ = 0.966) | logGEF = -0.558 * logW – 11.640 | 5.72 × 10^-13^ | 2.14 × 10^-13^ |
| DTX201 | logGEF = -0.722 * logW  – 10.839 (r^2^ = 0.899) | logGEF = -1.055 * logW – 11.202 | 1.33 × 10^-12^ | 7.11 × 10^-14^ |
| SPK-8011 | logGEF = -0.547 * logW  – 10.088 (r^2^ = 0.778) | logGEF = -0.960 * logW – 10.534 | 1.66 × 10^-11^ | 4.26 × 10^-13^ |

**Table S6. W^-0.25^-dependent allometric equations and human predictions for gene efficiency factor (GEF)**

| Vector | Allometric equation using W^-0.25^ | | Human GEF (molecule/day/vg for FIX) (IU/day/vg for FVIII) | |
| --- | --- | --- | --- | --- |
|  | Three species | Two species | Observed | Predicted |
| rAAV2-CMV-FIX | logGEF = 0.400 * W^-0.25^ + 1.635 (r^2^ = 0.989) | logGEF = 0.431 * W^-0.25^ + 1.560 | 66.8 | 51.1 |
| rAAV2-hAAT-FIX | logGEF = 0.533 * W^-0.25^ + 2.443 (r^2^ = 0.946) | logGEF = 0.444 * W^-0.25^ + 2.667 | 296 | 661 |
| scAAV2/8-LP1-hFIXco | logGEF = 1.461 * W^-0.25^ + 2.543 (r^2^ = 0.967) | logGEF = 1.264 * W^-0.25^ + 3.017 | 520 | 2745 |
| rAAV-Spark100-FIX-R338L (SPK-9001) | logGEF = 1.144 * W^-0.25^ + 2.311 (r^2^ = 0.984) | logGEF = 1.249 * W^-0.25^ + 2.054 | 737 | 296 |
| GO-8 | logGEF = 0.957 * W^-0.25^ – 11.189 (r^2^ = 0.999) | logGEF = 0.940 * W^-0.25^ – 11.148 | 1.30 × 10^-11^ | 1.50 × 10^-11^ |
| SB-525 | logGEF = 1.029 * W^-0.25^ – 11.879 (r^2^ = 0.884) | logGEF = 0.711 * W^-0.25^ – 11.128 | 1.19 × 10^-12^ | 1.31 × 10^-11^ |
| BMN270 | logGEF = 0.661 * W^-0.25^ – 12.445 (r^2^ = 0.999) | logGEF = 0.642 * W^-0.25^ – 12.400 | 5.72 × 10^-13^ | 6.64 × 10^-13^ |
| DTX201 | logGEF = 1.101 * W^-0.25^ – 12.365 (r^2^ = 0.992) | logGEF = 1.186 * W^-0.25^ – 12.565 | 1.33 × 10^-12^ | 7.00 × 10^-13^ |
| SPK-8011 | logGEF = 0.885 * W^-0.25^ – 11.317 (r^2^ = 0.939) | logGEF = 1.079 * W^-0.25^ – 11.774 | 1.66 × 10^-11^ | 3.84 × 10^-12^ |

**Table S7. Clinical immunogenicity data of eight vectors by dose cohort**

| Gene therapy | Route of administration | Dose (vg/kg) | Number of patients | Safety data | Refs |
| --- | --- | --- | --- | --- | --- |
| rAAV2-CMV-FIX | IM injection | 2x10^11^ | 3 | Increased anti-AAV antibody titers (n=8) and neutralizing antibodies (NABs) to AAV (n=7) | ^26^ |
|  |  | 6x10^11^ | 3 |  |  |
|  |  | 1.8x10^12^ | 2 |  |  |
| rAAV2-hAAT-FIX | IV infusion | 8x10^10^ | 2 | Increase in ALT and AST (n=2)  **Two patients in the high dose cohort developed T-cell response against capsid**. | ^15, 27^ |
|  |  | 4x10^11^ | 3 |  |  |
|  |  | 2x10^12^ | 2 |  |  |
| scAAV2/8-LP1-hFIXco | IV infusion | 2x10^11^ | 2 | Increased AAV8-IgG antibody titers (n=2) | ^28^ |
|  |  | 6x10^11^ | 2 | **T-cell response against capsid (n=2),** Increased AAV8-IgG antibody titers (n=2) |  |
|  |  | 2x10^12^ | 6 | **T-cell response against capsid (n=5),** elevation in ALT (n=4), Increased AAV8-IgG antibody titers (n=6) |  |
| SPK-9001 | IV infusion | 6x10^11^ | 10 | Increase in ALT (n=2) | ^17^ |
| AMT-060 (AAV5-FIX) | IV infusion | 5x10^12^ | 5 | Increase in ALT (n=1) | ^19^ |
|  |  | 2x10^13^ | 5 | Increase in ALT (n=3) |  |
| AMT-061 (AAV5-FIX Padua) | IV infusion | 2x10^13^ | 3 | Increase in ALT and AST (n=1) | ^20^ |
|  |  | 2x10^13^ | 31 | Transient transaminitis requiring  corticosteroids  (n=7) | ^21^ |
| GO-8 | IV infusion | 6x10^11^ | 1 | Increase in ALT (n=2) | ^29^ |
|  |  | 2x10^12^ | 2 |  |  |
| SB-525 |  | 9x10^11^ | 2 | Increase in ALT (n=4) | ^4^ |
|  |  | 2x10^12^ | 2 |  |  |
|  |  | 1x10^13^ | 2 |  |  |
|  |  | 3x10^13^ | 5 | Increase in ALT (n=4) |  |
| BMN270 | IV infusion | 6x10^12^ | 1 | Increase in ALT and anti-AAV5 antibodies (n=1) | ^8^ |
|  |  | 2x10^13^ | 1 | Anti-AAV5 antibodies(n=1) |  |
|  |  | 6x10^13^ | 7 | Increase in ALT and anti-AAV5 antibodies (n=7) |  |
| DTX201 | IV infusion | 5x10^12^ | 2 | N.A. | ^11^ |
|  |  | 1x10^13^ | 2 | Increase in ALT and AST (n=1) |  |
|  |  | 2x10^13^ | 4 | Increase in ALT and AST (n=2) |  |
| SPK-8011 | IV infusion | 5x10^11^ | 2 | None | ^13^ |
|  |  | 1x10^12^ | 3 | None |  |
|  |  | 1.5x10^12^ | 4 | Increase in ALT (n=4) |  |
|  |  | 2x10^12^ | 9 | Increase in ALT (n=3) |  |
